# A Multivalent Polyomavirus Vaccine Elicits Durable Neutralizing Antibody Responses in Macaques

**DOI:** 10.1101/2022.09.26.509096

**Authors:** Alberto Peretti, Diana G. Scorpio, Wing-Pui Kong, Yuk-Ying S. Pang, Michael McCarthy, Kuishu Ren, Moriah Jackson, Barney S. Graham, Christopher B. Buck, Patrick M. McTamney, Diana V. Pastrana

## Abstract

In 2019, there were about 100,000 kidney transplants globally, with more than a quarter of them performed in the United States. Unfortunately, some engrafted organs are lost to polyomavirus-associated nephropathy (PyVAN) caused by BK and JC viruses (BKPyV and JCPyV). Transplant patients are routinely monitored for BKPyV viremia, which is an accepted hallmark of nascent nephropathy. If viremia is detected, a reduction in immunosuppressive therapy is standard care, but the intervention comes with increased risk of immune rejection of the engrafted organ. Recent reports have suggested that transplant recipients with high levels of polyomavirus-neutralizing antibodies are protected against PyVAN. Virus-like particle (VLP) vaccines, similar to approved human papillomaviruses vaccines, have an excellent safety record and are known to induce high levels of neutralizing antibodies associated and long-lasting protection from infection. In this study, we demonstrate that VLPs representing BKPyV genotypes I, II, and IV, as well as JCPyV genotype 2 produced in insect cells elicit robust antibody titers. In rhesus macaques, all monkeys developed neutralizing antibody titers above a previously proposed protective threshold of 10,000. A second inoculation, administered 19 weeks after priming, boosted titers to a plateau of ≥25,000 that was maintained for almost two years. No vaccine-related adverse events were observed in any macaques. A multivalent BK/JC VLP immunogen did not show inferiority compared to the single-genotype VLP immunogens. Considering these encouraging results, we believe a clinical trial administering the multivalent VLP vaccine in patients waiting to receive a kidney transplant is warranted to evaluate its ability to reduce or eliminate PyVAN.

**HIGHLIGHTS:**

- Recombinant virus-like particle vaccine was safely administered to rhesus macaques
- Vaccination generated high-titer neutralizing antibody responses
- Multivalent BK/JC polyomavirus vaccine was as effective as monovalent vaccines
- High neutralizing titers were sustained for 92 weeks without appreciable decline

## INTRODUCTION

When kidney transplantation was first being established, up to 90% of cases ended in acute immunological rejection of the allograft (reviewed in [1]). The use of immunosuppressive treatments such as calcineurin inhibitors (cyclosporine, tacrolimus), corticosteroids, T-cell depleting antibodies, interleukin 2 receptor antagonists, etc. dramatically improved the outcomes. An unforeseen consequence of this success was an increase in polyomavirus associated nephropathy (PyVAN), which can occur in up to 10% of kidney transplant recipients (KTRs) [2–6]. PyVAN is also a significant risk for other types or solid organ transplant patients[7–9].

BK polyomavirus (BKPyV) and JC polyomavirus (JCPyV) are closely related species that establish chronic infections in nearly all humans [10, 11]. BKPyV consists of four genotypes, numbered I through IV, with several sub-genotypes. The four genotypes are distinct serotypes, meaning that antibody responses capable of neutralizing the infectivity of one genotype do not necessarily neutralize other genotypes [12, 13]. JCPyV consists of seven genotypes [14]. Although the different JCPyV genotypes are closely related and are not generally considered distinct serotypes, some individuals have neutralizing “blind spots” that JCPyV variants can occupy after accumulating point mutations [15]. These viruses are believed to maintain a latent infection in their hosts, with periodic reactivation in healthy individuals. Reactivation appears to be more common when immunity is weakened, such as during pregnancy [16] or in old age [17]. BKPyV is the more common culprit in PyVAN, with JCPyV being responsible for only a few percent of cases [18, 19].

Kidney transplant recipients (KTRs) are routinely monitored for the appearance of BKPyV viremia and rising creatinine, which presage PyVAN. If BKPyV is detected above a threshold of 10,000 copies per ml of blood, standard care is to decrease the level of immunosuppressive therapy, with the aim of bringing the polyomavirus infection under control [20]. However, reducing or switching immunosuppressants carries an inherent risk of precipitating immune rejection of the transplanted organ [21, 22].

In the absence of specific antivirals for BKPyV, some researchers have used adjunctive therapies such as intravenous immunoglobulin (IVIG) [23–25] or drugs with broad anti-viral activity, such as cidofovir [26–28], leflunomide [29–31] or ciprofloxacin [32, 33] in an attempt to treat the infection, but their usefulness remains inconclusive.

PyVAN was traditionally thought to arise mainly from reactivation of latent infection in the kidney transplant recipient. More recent evidence supports the idea of a donor-derived infection that “hitchhikes” in the engrafted organ. All of the following circumstances increase the probability of developing PyVAN: 1) differences in serostatus between the kidney donor and recipient [34–36], 2) presence of polyomavirus viruria in the donor around the time of transplantation [37, 38], and 3) correlation of donor but not recipient genotype replication [37, 39–41]. Additional evidence comes from isolated case reports where PyVAN genotype is identical in two recipients who received kidneys from the same deceased allograft donor [34, 42].

Although natural infection with BKPyV and JCPyV early in life can result in high titer neutralizing antibody responses, most individuals do not cross-neutralize all variants of the two virus species [15, 43]. KTRs who have reciprocal EC_50_ neutralizing antibody titers >10,000 against BKPyV genotypes present in the kidney donor are unlikely to develop viremia or PyVAN after transplantation [40]. It thus stands to reason that a vaccine capable of inducing neutralizing titers >10,000 against all BKPyV and JCPyV genotypes may protect transplant patients against PyVAN. Such a vaccine could be administered while the patient is on the organ transplant wait list, such that high levels of neutralizing antibodies will be present at the time of transplant.

Polyomaviruses are non-enveloped icosahedral viruses. The major capsid protein, VP1, can spontaneously self-assemble into virus-like-particles (VLP) that closely resemble the surface of native polyomavirus virions. Results for VLP vaccines targeting a related virus family, human papillomaviruses (HPVs), have shown that rigidly repetitive structures can induce exceptionally potent and durable neutralizing antibody responses [44]. BKPyV and/or JCVPyV VLPs have been successfully produced in E.coli [45], mammal [12], baculovirus [46–49] and yeast [50, 51] based expression systems. Preclinical studies using VLPs representing human polyomaviruses to immunize mice or rabbits show that they produce high titer neutralizing antibody responses [12, 13, 46, 52, 53]. In this study, we show that immunization of monkeys with a multivalent PyV-VLP immunogen produced using a standard baculovirus-based expression system induced sustained neutralizing antibody responses well above the previously established correlate of protection against PyVAN.

## MATERIALS AND METHODS

### VLP production

The genes encoding for VP1 from BKPyV and JCPyV were codon modified for use in insect cells and synthesized by ThermoFisher Scientific (Waltham, MA). BKPyV VP1 genes were from three different genotypes: BKPyV-Ib2 isolate PittVR2 (GenBank # DQ989796); BKPyV-II isolate GBR-12 (GenBank # AB263920); and BKPyV-IVc2 isolate A-66H (GenBank # AB369093). One JCPyV genotype was used: JCPyV-5053 (genotype 2A) [15]. The synthetic genes were cloned into the BamHI and HindIII restriction enzyme sites of the baculovirus expression vector pFastBac1 (ThermoFisher). Per manufacturer’s instructions, the recombinant pFastBac plasmids were transformed into E. coli cells containing a baculovirus shuttle vector and helper plasmid (DH10Bac cells; ThermoFisher) to obtain colonies containing recombinant bacmids. The bacmids were purified and used to transfect Sf9 cells (ThermoFisher) to obtain a recombinant baculovirus stock that was subsequently amplified and titered.

One liter of High 5 cells at a density of 2.2 million cells/ml were infected at a multiplicity of infection of one with the recombinant baculovirus stocks. After 3 days, the infected cells were harvested by centrifugation at 1000 x g for 15 minutes. The clarified supernatant was retained. The cell pellet was resuspended in one volume of DPBS with calcium and magnesium (ThermoFisher). The resuspended pellet and clarified primary supernatant were processed in parallel as follows: added MgCl_2_ to 10 mM final concentration, benzonase to 0.1% (v/v), Triton-X100 to 0.5% v/v (all from MilliporeSigma, St. Louis, MO) and incubated at 37°C for 1.5hrs. Samples were frozen until ready to use.

Purification of the polyomavirus VLPs was performed similarly to VLP purification from mammalian cells [12] https://ccrod.cancer.gov/confluence/display/LCOTF/PseudovirusProduction. Briefly, the pellets were thawed; cOmplete™ EDTA-free protease inhibitor cocktail (MilliporeSigma) and 1M ammonium sulfate pH 9 were added to adjust the pH to 7 and the mixture was incubated at 37°C for 1hr. NaCl was added to a final concentration of 800 mM and the lysate was clarified at 12,000 x g for 10 minutes. The pellet was resuspended in 700 μl of DPBS with 800 mM NaCl and re-clarified and a secondary supernatant was saved. The secondary supernatants were combined and loaded onto a 27, 33 and 39% iodixanol (Optiprep from MilliporeSigma) step gradient. The gradient was centrifuged at 234,000 x g for 3.5 hrs and fractions collected. Fractions were screened by Quant-it Picogreen dsDNA reagent and Pierce BCA protein assay (both from ThermoScientific). Fractions suspected of containing VLPs were analyzed using negative stain electron microscopy and staining of SDS-PAGE gels. VLP containing fractions were further purified with two rounds of size exclusion chromatography through a Sephacryl S-400 column (Cytvia, Marlborough, MA) and analyzed by multi-light scattering to choose relevant fractions. The material was concentrated to 1 mg/ml with a centricon unit (MilliporeSigma).

For some cultures, it appeared that spontaneous cell lysis released substantial amounts of VLPs into the culture medium. VLPs were recovered from clarified supernatants by pelleting the VLPs onto an Optiprep cushion in 30 ml tubes. Iodixanol was removed from particles using a Superose 6 column (Cytvia). Particles were then concentrated and further purified with a Sephacryl S-400 column before finally being concentrated to approximately 1 mg/mL in PBS.

The resulting purified VLPs from the primary supernatant (sVLPs) and pellets (pVLPs) were quantified by four different methods: Pierce BCA protein assay, absorbance at 280 nm in a spectrophotometer, Quant-it protein assay kit (ThermoFisher), and Syproruby protein gel stain (ThermoFisher).

### Immunizations

Mouse and macaque experiments were conducted in accordance with MedImmune’s Institutional Animal Care and Use Committee (IACUC) and the Vaccine Research Center/National Institutes of Health’s IACUC. Animals were housed in accordance with the Guide for the Care and Use of Laboratory animals, Eight Edition, 2001, National Research Council.

Five female BALB/c mice per group were immunized intramuscularly with 1 μg of purified VLPs adjuvanted with 100 μg of Alhydrogel (Brenntag) by mixing. The mice received a second immunization four weeks after the priming dose. Blood samples were taken from the mandibular vein before immunization, and two weeks after the first and second immunizations. For the sVLPs there were five groups: 1) BKPyV-I; 2) BKPyV-II; 3) BKPyV-IV; 4) JCPyV; 5) combination of the three BKPyV genotypes plus JCPyV. Group 5 received a total of 4 μg of VLPs (one microgram per genotype). For the pVLPs some of the JCPyV VLP material was insufficient so only 4 groups were immunized: 1) BKPyV-I; 2) BKPyV-II; 3) BKPyV-IV; 4) combination of the three BKPyV genotypes.

Nine adult Rhesus macaques with ages ranging from 16 to 20 years were immunized intramuscularly (left quadricep) in three groups: 1) 10 μg BKPyV-I pVLPs; 2) 10 μg BKPyV-IV pVLPs; and 3) a mixture of 10 μg each of BKPyV-I, BKPyV-II, BKPyV-IV pVLPs and JCPyV sVLPs. VLPs were adjuvanted with 100 μg Alhydrogel (Brenntag) by mixing. The monkeys received a second immunization nineteen weeks after the prime. Blood was collected via femoral vein prior to immunization, weekly until six weeks post prime, and then at weeks 25, 43 and 92.

### Neutralization Assays

BKPyV, JCPyV and SV40 polyomavirus neutralization assays were performed using 293TT derived pseudovirions containing NanoLuc luciferase reporter plasmids pcsNuc and phsNuc [54] as described previously [13, 15]. Briefly, 293TT cells were plated at a density of 30,000 cells per well in 96-well tissue culture treated plates. Pseudovirions were incubated with 5-fold serial dilutions of the mouse or primate sera for 1 hour at 4°C in polystyrene untreated 96 well plates. The pseudovirus/serum mixture was then added to pre-plated cells and returned to an incubator for 72 hours. Twenty-five microliters of conditioned media were transferred to white luminometry plates (Perkin-Elmer) and used to evaluate the presence of secreted NanoLuc. The Nano-Glo Luciferase Assay system (Promega) working substrate was diluted 1:8 in PBS and 80 μl was added to the conditioned media. Signal was detected with a POLARstar optima luminometer (BMG), with gain ranging from 2,200 to 2,800 depending on the viral stock. The EC_50_ (reciprocal 50% neutralization dilution) was determined using Prism software (GraphPad) to calculate the best fit for a variable-slope sigmoidal dose-response curve.

## RESULTS

Baculovirus-based expression of BKPyV and JCPyV VP1 proteins has been shown to generate high-quality virus-like particles (VLPs)[46–49]. Our experiments using a standard commercial baculoviral vector system confirmed these observations. For BKPyV (depending on genotype), we were able to obtain 6-30 mg/L of VLPs from insect cells seeded at a density of 2.2 million cells/ml; for JCPyV, we were able to obtain ~0.6 mg/L. The final VLP preparations had a high level of purity, with full-length VP1 accounting for a large majority of the protein signal in an SDS-PAGE analysis (Fig. 1A). VLPs obtained from the cellular pellet (pVLPs) or supernatants (sVLPs) of the baculovirus-infected cultures had similar levels of purity. VLPs are stabilized by intermolecular disulfide bonds between VP1 monomers, as evidenced by non-reducing SDS-PAGE analysis, which shows slower-migrating bands corresponding to disulfide-linked VP1 multimers (Supplementary Figure 1). In addition, electron microscopic analysis showed homogeneous particles of the expected ~50 nm diameter (Fig 1B to 1E).

**Figure 1.**
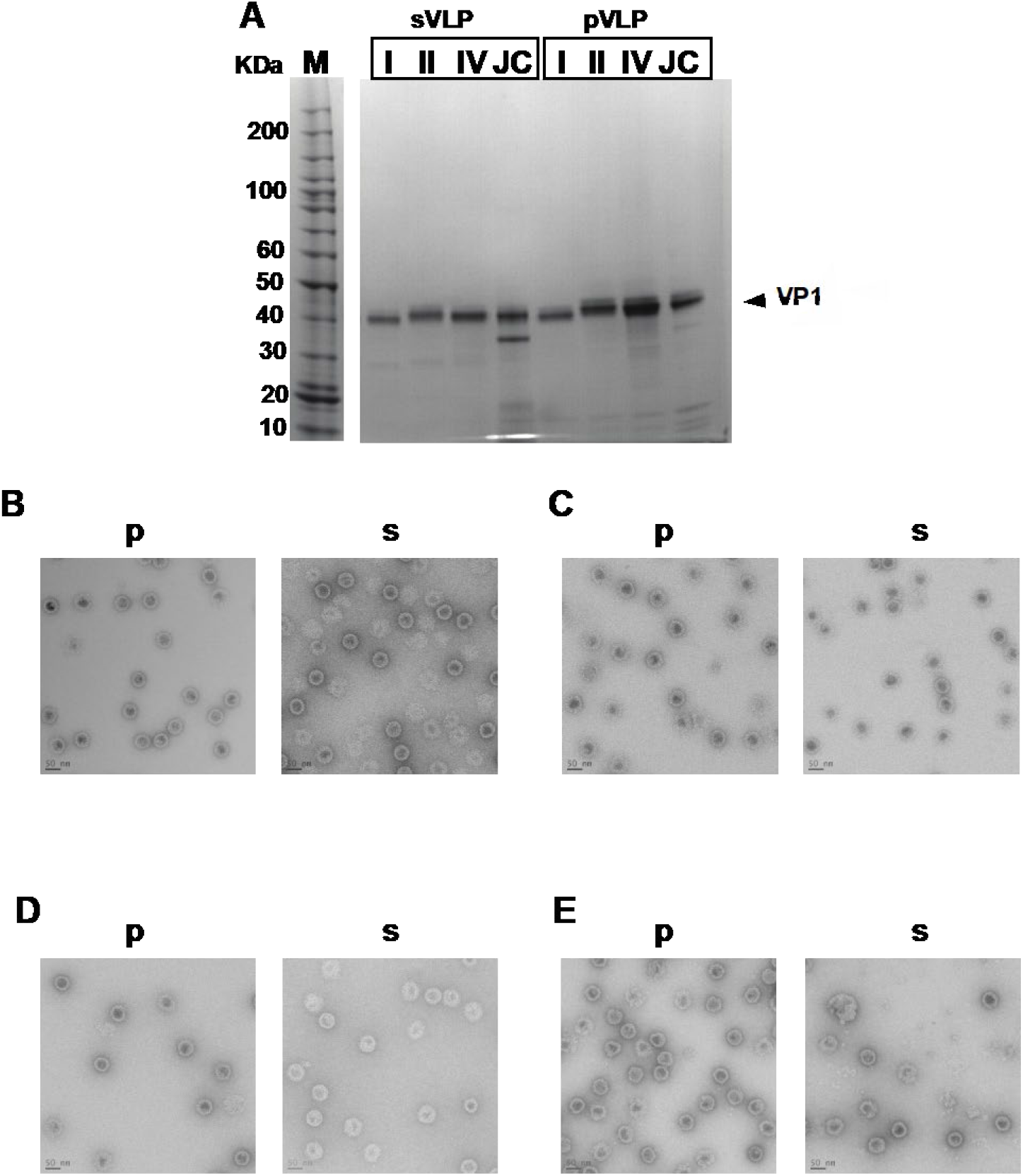
Production and purification of polyomavirus VLPs: A) Coomassie-stained SDS-PAGE analysis of VLPs used to immunize mice and monkeys. Lanes 1-4 correspond to culture supernatant-derived VLPs (s) and lanes 5-8 correspond to cellular pellet-derived VLPs (p). Lanes 1,5 BKPyV-I (I), lanes 2,6 BKPyV-II (II), lanes 3,7 BKPyV-IV (IV), and lanes 4,8 JCPyV (JC). B-E) Electron micrographs of purified indicated VLPs. A bar corresponding to 50nm in length was added for size comparisons.

VLPs produced in mammalian cells have been shown to generate high-titer cognate anti-BKPyV and JCPyV neutralizing responses in mice [13, 15]. The baculovirus-based VLPs showed a similar ability to generate high-titer neutralizing antibody responses, with a reciprocal log_10_ 50% neutralizing dilution (EC_50_) titer of at least 3.9 (BKPyV-IV) and as high as 4.9 (BKPyV-II) (Figure 2). BKPyV antibody titers from primed mice were absent or more than 10-fold lower against non-cognate types, except for BKPyV-IV-primed mice, which cross-neutralized BKVPyV-II. None of the BKPyV-primed mice showed detectable JCPyV-neutralizing antibody responses, but JCPyV-primed mice showed low-level cross-neutralization of all BKPyV genotypes. Multivalent priming of mice with a mixture of all three BKPyV types plus JCPyV generated neutralizing titers comparable to the titers obtained when a monovalent immunogen was used. A second injection with a monovalent BKPyV vaccine boosted cognate titers and induced robustly cross-neutralizing responses against non-cognate BKPyV genotypes. Sera from BKPyV immunized mice were still unable to neutralize JCPyV pseudovirions, with the exception of mice immunized twice with BKPyV-II sVLPs, which neutralized JCPyV pseudovirions with an EC_50_ titer of 3.6. Two immunizations with the multivalent vaccine generated high neutralizing titers against all BKPyV types and JCPyV.

**Figure 2.**
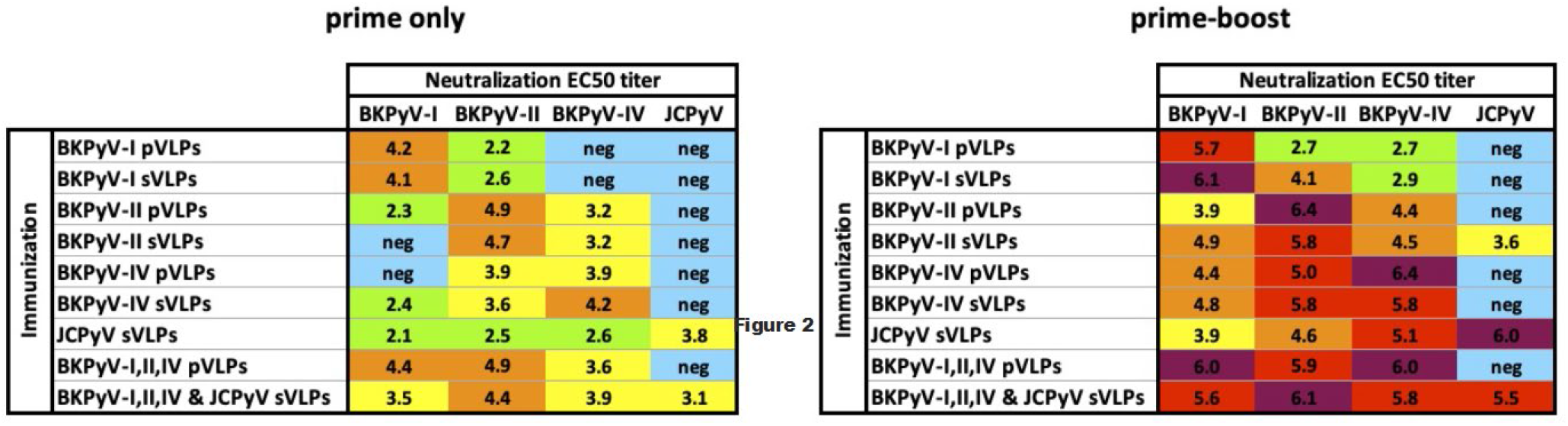
Serological response of mice immunized with polyomavirus VLPs: Pseudovirus neutralization serology was performed on pooled sera from groups of five BALB/c mice after the first (prime) and second (boost) doses of VLPs administered intramuscularly with alum adjuvant. EC_50_ titers of 6 and above are highlighted in purple, 5 to 5.9 in red, 4 to 4.9 in orange, 3 to 3.9 in yellow, 2 to 2.9 in green, and negative (undetectable to 1.9) in blue.

Captive Rhesus macaques often harbor clinically inapparent infections with SV40, which is closely related to BKPyV and JCPyV [55]. The presence of anti-SV40 neutralizing antibodies prior to human polyomavirus VLP inoculation was therefore investigated in monkeys selected for the present study. All nine monkeys had high-titer neutralizing antibodies specific for SV40 at study entry, with titers ranging from 4.6 to 6.2 (Table S1). These pre-immunization sera were also investigated for cross-reactivity to human polyomaviruses. A variety of previously reported BKPyV and JCPyV variants were tested [13, 15, 56–59]. Each of the macaques had weak or moderately cross-neutralizing serum antibodies against some BKPyV or JCPyV variants, consistent with older literature[60].

Six days after immunization with a single dose of VLPs all animals showed a dramatic increase in neutralizing titers against cognate and heterologous BKPyV and JCPyV genotypes (Figure 3). Although some monkeys that received only BKPyV-I or BKPyV-IV VLPs showed a major an increase in anti-JCPyV titers, the magnitude of the increase was not uniform. Immunization with a mixture of BKPyV and JCPyV VLPs resulted in high neutralizing titers even after just a priming dose, with log EC_50_s of 2.8 to 6.0. Macaques were boosted on week 19 with the same immunogen/s they received in the initial inoculation. Pathogenesis-associated mutations – such as BKPyV E73Q and JCPyV L55F, and S269F - were neutralized at similar titers to the wild type strains (Table S1). The booster dose increased the neutralizing titers of the macaques by less than 1 log, except for one macaque immunized with the combination of BKPyV and JCPyV that achieved an improvement of 1.3 logs in titer against JCPyV.

**Figure 3.**
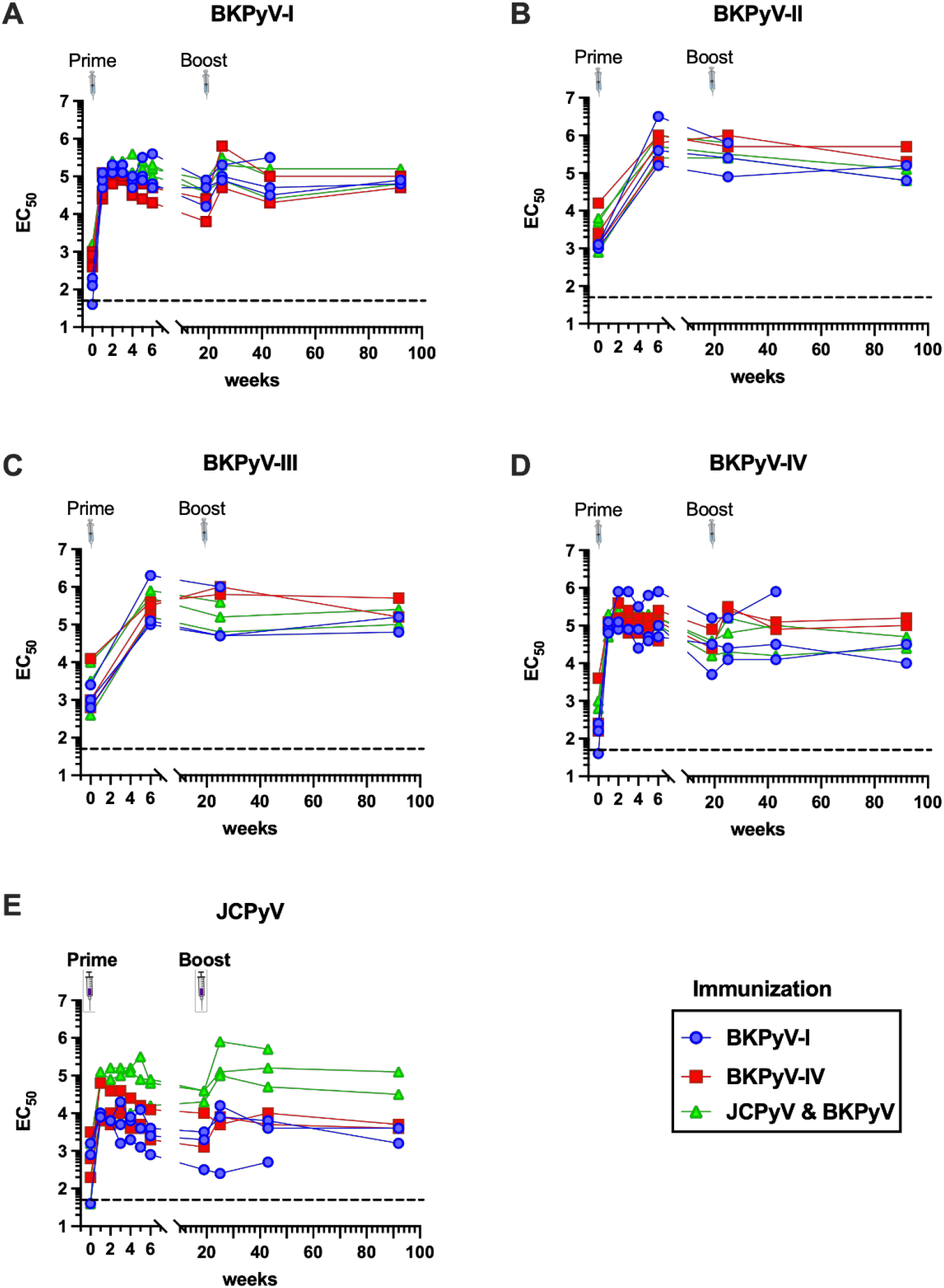
Serological response of rhesus macaques immunized with polyomavirus VLPs: 16- to 20-year-old macaques were immunized with 10 μg of VLPs and 100 μg of alum were used as an adjuvant. Serum samples were obtained prior to immunization and for the first 6 weeks after administration of the initial dose of VLPs. A second dose of VLPs was given at week 19 post prime. Additional serum was obtained at weeks 25, 43 and 92. The neutralizing titer of the sera was evaluated over time and the EC_50_ is depicted in the graphs. Macaques immunized with BKPyV-I are depicted by the blue circles, those immunized with BKPyV-IV are shown with red squares, and macaques receiving a quadrivalent vaccine (BKPyV-I, II, IV and JCPyV) are shown with green triangles.

The neutralizing antibody response of macaques was followed for almost two years. One animal from each of the immunization groups died during the study. These deaths were unlikely the result of the vaccine, but rather due to older age. The average lifespan of rhesus macaques in captivity is approximately 25 years [61], and the average age of the monkeys at study entry was 18 years of age. Necropsies showed that one animal from the BKV-IV immunized group died at age 22 from a suspect intestinal adenocarcinoma (which is commonly observed in aged macaques) 7 weeks after the priming dose. An animal from the BKV-I immunized group was euthanized 34 weeks post-boost due to old age and lethargy. A third animal was euthanized 1 year and 1 month after the booster immunization due to organ failure from a suspected heart attack or stroke. The neutralizing titers of surviving macaques remained stable for at least 92 weeks, the last time point that was analyzed (Figure 3). The titers remained high during the entire course of the study, with the lowest homologous EC_50_ titer being 4.4 against BKPyV-IV from a macaque immunized with the BKPyV and JCPyV mixture. This titer is above the 4.0 level shown to correlate with protection against post-transplant PyVAN [40]. Remarkably, the EC_50_ titers also stayed above log 4.0 for BKPyV-I/IV cross-neutralizing responses for the duration of the study. For JCPyV, titers were also stable even for monkeys immunized with only BKPyV-I or -IV.

## DISCUSSION

VLPs present a potential vaccine platform to address post-transplant PyVAN, and vaccination studies in mice and particularly NHPs may provide models that translate to humans. Although mice were unable to elicit potent BKPyV cross-neutralizing Abs after priming, once the mice were boosted with a second immunization, they were able to generate a more broadly neutralizing response against all heterologous genotypes. In macaques, priming alone resulted in a broadly cross-neutralizing heterologous response. The macaques had strong pre-existing humoral responses to SV40, which we show elicits low or moderate levels of BKPyV and JCPyV cross-neutralizing antibody responses. The first immunization in the monkeys could thus be thought of as functionally a booster dose. Indeed, there is some evidence that infection with one of the nephropathic polyomaviruses (BKPyV and JCPyV) elicits some level of T cell-mediated protection against infection with the other species [62]. Additionally, the possibility that the macaques have been exposed to reverse zoonosis with BKPyV and JCPyV cannot be eliminated, as the animals are in close contact with their human handlers. The observation that the vaccine is successful in boosting and broadening the pre-existing immune responses in macaques is encouraging, as it roughly reflects the situation for candidate organ transplant recipients, who are typically already infected with one or more BKPyV or JCPyV genotypes. The pre-existing neutralizing antibody responses in the monkeys at study entry did not always correlate with how much the immune response improved after vaccination with VLPs, particularly after the first dose. For example, the pre-vaccination titer of one macaque that had a reciprocal log EC_50_ titer of 3.3 against BKPyV genotype IV (with a 73Q mutation) had the same post immune titer (4.6) as a macaque that had no detectable pre-vaccination neutralizing antibodies. This is an argument for having a two-dose multivalent vaccine that will ensure the most diverse response against all genotypes and variants.

In the context of KTRs, uncontrolled polyomavirus infection allows for the evolution of polyomaviruses that accumulate neutralization escape mutations in the VP1 gene [56, 63]. Some of these mutations are concentrated in the VP1 loop associated with receptor binding (BC-loop). Among these is the E73K mutation, which substitutes a negatively charged amino acid side chain for a positively charged one. The neutralizing titers against this mutant were the lowest in both genotype I and IV backgrounds, in some instances falling below the presumed protective EC_50_ of 4.0. Vaccination of patients on organ transplant waitlists could set up a scenario where replication of wild-type BKPyV strains is blocked, such that pathogenesis-associated mutations, such as E73K, are unable to accumulate. In the non-human primates, the antibody responses neutralized not only cognate types, but also pathogenesis-associated mutations.

Patients with end stage kidney disease face many challenges. After their kidneys fail, they might receive dialysis treatments, a debilitating process that consumes 3-5 hours three times per week. A person might remain on dialysis an average of 3-5 years before a kidney becomes available. In addition to the morbidity associated with kidney disease, the cost of a kidney transplant was estimated to be around $442,500 in the year 2020 [64]. It is imperative to preserve the health of engrafted organs. As discussed in the introduction, compounds with antiviral activity, use of IVIG, and lowering/switching immunosuppressive drugs to stave off PyV infections and preserve kidney function, all have limited effectiveness. A prophylactic vaccine approach might prove more effective in the fight against loss of kidneys due to PyVAN or to immune rejection following the reduced immunosuppressive therapy nascent PyVAN necessitates. VLP based vaccines have an excellent safety record and tend to elicit high titer durable responses. Waiting lists to obtain a donated organ produce an excellent window of opportunity during which VLP based vaccine doses could be administered. Our data suggest that vaccination could durably arm patients with high-titer neutralizing antibodies against all BKPyV and JCPyV variants.

Current clinical trials seek to evaluate the safety and efficacy of preventing viremia with BKPyV-neutralizing monoclonal antibodies (mAbs) or pooled antibody preparations administered intravenously (Clinical trial identifiers NCT04294472, NCT05264259, NCT04222023 and NCT05358106). If the results of these trials are favorable, they will support the idea that a vaccine that is able to generate high titer neutralizing antibodies would also be effective, particularly if given before immunosuppressives are administered. The vaccine could be less expensive, easier to administer, and more durable than mAb therapeutics. BK and JC polyomaviruses are also known to contribute to hemorrhagic cystitis in bone marrow transplant recipients.

Recent evidence suggests that polyomaviruses contribute to the increased incidence of bladder and kidney cancer in organ transplant patients [65]. An added benefit of an anti-polyomavirus vaccine could be a decrease in the risk of urothelial neoplasias, which organ transplant recipients are more prone to developing. These polyomaviruses are being increasingly recognized as having deleterious effects in the kidneys of other solid organ transplants such as liver, heart and lung [7–9]. The BKPyV and JCPyV vaccine should be considered as a possible preventive intervention for organ transplant recipients. Moreover, the development of the first human polyomavirus vaccine should pave the way for developing vaccines against other disease-causing polyomaviruses.

## ABBREVIATIONS

VLP: Virus-like particle
sVLP & pVLP: supernatant or pellet derived VLP
PyVAN: Polyomavirus associated nephropathy
KTR: Kidney transplant recipient

## AUTHOR CONTRIBUTIONS

Alberto Peretti: conceptualization, acquisition and analysis of data, Writing – review and editing

Diana G. Scorpio: conceptualization, acquisition and analysis of data, Writing – review and editing

Wing-Pui Kong: acquisition, writing – review and editing

Yuk-Ying S. Pang: acquisition and analysis of data, Writing – review and editing Michael McCarthy: acquisition and analysis of data

Kuishu Ren: acquisition and analysis of data

Moriah Jackson: acquisition and analysis of data

Barney S. Graham: conceptualization, provision of resources, editing

Patrick McTamney, conceptualization, acquisition and analysis of data, Writing – review and editing

Christopher B. Buck, conceptualization, analysis of data, Writing – review and editing Diana V. Pastrana, conceptualization, acquisition and analysis of data, Writing – original draft

## ACKNOWLEDGEMENTS

This Research was supported by the Intramural Research Program of the National Cancer Institute and the National Institute of Allergy and Infectious Diseases. We are grateful to Elizabeth McCarthy for her aid in scheduling the study and supporting logistics.

