## Supplemental Figure 1 for "A Multivalent Polyomavirus Vaccine Elicits Durable Neutralizing Antibody Responses in Macaques"

### Supplementary Figure 1: Non-reducing gel analysis of VP1 disulfide cross-linking

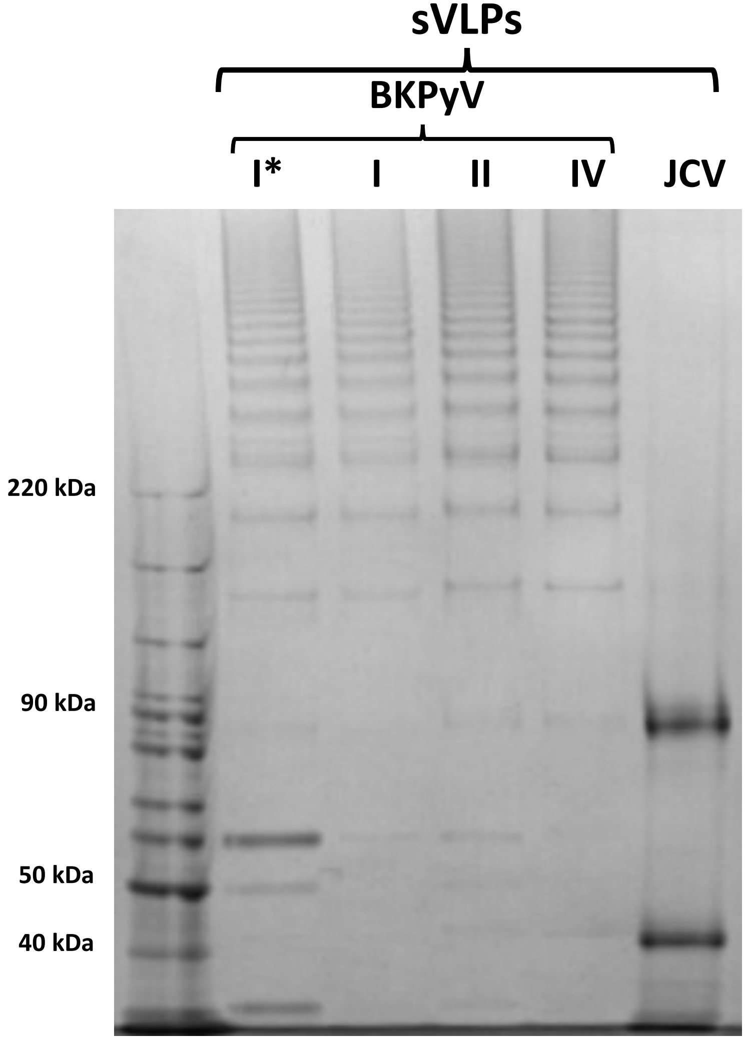


*this BKPyV-I preparation was purified without Triton or benzonase at the time of harvest

A protocol for performing non-reducing gel analysis on VLPs con be found on:
https://ccrod.cancer.gov/confluence/display/LCOTF/ImprovedMaturation
